# THE ROLE OF POTASSIUM CHANNELS IN THE PATHOGENESIS OF GASTROINTESTINAL CANCERS AND THERAPEUTIC POTENTIAL

**DOI:** 10.1101/2020.03.10.984039

**Authors:** David Shorthouse, John L Zhuang, Eric P Rahrmann, Cassandra Kosmidou, Katherine Wickham Rahrmann, Michael Hall, Benedict Greenwood, Ginny Devonshire, Richard J Gilbertson, Rebecca C. Fitzgerald, Benjamin A Hall

## Abstract

Voltage sensitive potassium channels play an important role in controlling membrane potential and ionic homeostasis in the gut and have been implicated in gastrointestinal (GI) cancers. Through large scale analysis of 1594 patients with GI cancers coupled with *in vitro* models we find KCNQ family genes are mutated in ~30% of patients, and play therapeutically targetable roles in GI cancer growth. KCNQ1 and KCNQ3 mediate the WNT pathway and MYC to increase proliferation, and its resultant effects on cadherins junctions. This also highlights novel roles for KCNQ3 in non-excitable tissues. We additionally discover that activity of KCNQ3 sensitises cancer cells to existing potassium channel inhibitors, and that inhibition of KCNQ activity reduces proliferation of GI cancer. These findings reveal a novel and exploitable role for potassium channels in the advancement of human cancer, and highlight that supplemental treatments for GI cancers may exist through KCNQ inhibitors.

**SIGNIFICANCE:** KCNQ channels modulate the WNT pathway and MYC signalling, and drive growth of gastrointestinal cancers. Available drugs modulate these pathways and offer therapeutic potential in gastrointestinal cancer.

## INTRODUCTION

The *KCNQ* (Potassium Voltage-Gated Channel Subfamily Q) family of ion channels encode potassium transporters(1). KCNQ proteins typically repolarise the plasma membrane of a cell after depolarisation by allowing the export of potassium ions, and are therefore involved in wide ranging biological functions including cardiac action potentials(2), neural excitability(3), and ionic homeostasis in the gastrointestinal tract(4). Diseases resulting from loss or gain-of-function (LoF/GoF) mutations in the *KCNQ* family are also wide ranging, and include epilepsy(5), cardiac long and short QT syndrome(6), and Autism-like disorders(7). Due to their involvement in human disease, numerous molecules that interact with them are therapeutics. KCNQ1 interacts with a family of KCNE ancillary proteins in varying tissues, but is otherwise homotetrameric(1). KCNQ2, 3, 4, and 5 however, can interact with each other and the KCNE family to theoretically form hundreds of combinations of channels, but are predominantly found in KCNQ2/KCNQ3 heteromers in the brain.

There is preliminary evidence to suggest that members of the *KCNQ* family may contribute to the cancer phenotype. KCNQ1 plays a role in colon cancer(8), as well as in hepatocellular carcinoma(9), and KCNQ3 is hypermutated in oesophageal adenocarcinoma(10). Furthermore, we have previously identified that *KCNQ1* and *KCNQ3* RNA expression correlates with a cancer gene expression profile(11). These all hint to an involvement of *KCNQ* genes in the pathogenesis of gastrointestinal (GI) cancers. This might be expected since membrane transport is critical to the homeostatic function of luminal epithelial cells, but so far this has not been extensively explored, aside from a reported interaction between KCNQ1 and beta-catenin in the colorectal epithelium(8), which has not been observed in other tissues and has no clear link to clinical outcomes.

In this study, we investigate the mechanistic roles and therapeutic potential of the *KCNQ* family in gastrointestinal cancer by combining the study of highly annotated clinical and sequencing data sets of large numbers of patients (n = 1594) with *in vitro* cell culture assays on relevant cell lines. We find that KCNQ activity directly controls cancer cell growth through activating beta-catenin and MYC via the modulation of cadherins junctions, and that already clinically available drugs that interact with KCNQ channels are a promising therapeutic avenue for GI cancer.

## RESULTS

### *KCNQ* genes are highly altered in GI cancers

To fully characterise how *KCNQ/KCNE* genes are altered in GI cancer we studied all genetic alterations in the GI cohorts of The Cancer Genome Atlas (TCGA), combined with our own Oesophageal Adenocarcinoma data (OCCAMS) as part of the International Cancer Genome Consortium (ICGC), in which *KCNQ3* is recurrently missense mutated in 9.4% of patients(10). Cohorts were: Oesophageal Squamous Cell Carcinoma (OSCC, n = 103), Oesophageal Adenocarcinoma in two groups: the TCGA (n = 93)(12) and a subset of our own data for which full genetic analysis has been performed (n = 378)(10); Stomach Adenocarcinoma (STAD, n = 426)(13), and Colorectal Adenocarcinoma (COADREAD, n = 594)(14). 31% of all patients with GI cancers (n = 1594) had genetic alterations in at least one member of the *KCNQ/E* families.

From this dataset, we took several orthogonal approaches to assess the role of the KCNQ/E family in the cancer. We calculated the genetic status of all members of the KCNQ and modulatory KCNE gene families, as well several known driver genes in GI cancers (**Figure 1A**). We find a large number of amplifications of *KCNQ2* and *KCNQ3* (defined as copy number > 2 times the average ploidy). Both genes are in chromosomal regions commonly amplified in GI cancers (*KCNQ2*: chromosome 20q13.3, *KCNQ3*: chromosome 8q24.22) and known to be involved in cancer progression(15,16). *KCNQ3* in particular is located in a locus known to contain a large number of oncogenic protein coding and lncRNA genes (**Figure 1B**), including *MYC*, and is significantly (adjusted p <0.0001) co-associated with *MYC* amplifications (**Table S1**), thus many patients amplifying *MYC* will also amplify *KCNQ3*. Overall 174 (11%) of patients have a mutation/copy number change in *KCNQ3*, and whilst the 8q24 locus is a known susceptibility indicator in many cancers, this gene has not previously been explored in cancer. We additionally find that most alterations in *KCNQ1*, a gene already implicated as a tumor suppressor(8), are deletions or missense/truncating mutation events, indicating that this proposed role may extend beyond just colorectal adenocarcinoma where it has been previously studied. We also find several, significant (adjusted p < 0.05), mutually exclusive alteration events within the *KCNQ* family (**Table S1**), notably between *KCNQ1* and *KCNQ3* (adjusted p = 0.0003), and between *KCNQ2* and *KCNQ3, KCNQ4*, and *KCNQ5*. This pattern reveals that genetic alteration events generally occur in only a single *KCNQ* gene, so alteration to a single member may be sufficient to confer a selective advantage. Studying patient stage, we find no observed correlation between mutations in the *KCNQ/E* family and American Joint Committee on Cancer (AJCC) stage where annotated (**Figure S1A**). At the individual cancer level (**Figure S1B**), we find that with the exception of OSCC, which is primarily copy number driven(17), all cancers have an equal ratio of mutations and copy number changes, and that no single disease contains a majority of alterations. To identify functional significance of mutations in our cohort we also performed dN/dS analysis (18) (**Figure S1C**). dN/dS ratios show that, across all patients (n=1594), *KCNQ1* missense mutations are less common than expected (dN/dS < 1, q < 0.05), whilst KCNQ3 and KCNQ5 are under positive selection i.e. more common than expected (dN/dS > 1, q < 0.05) in OAC and STAD cohorts respectively.

**Figure 1:**
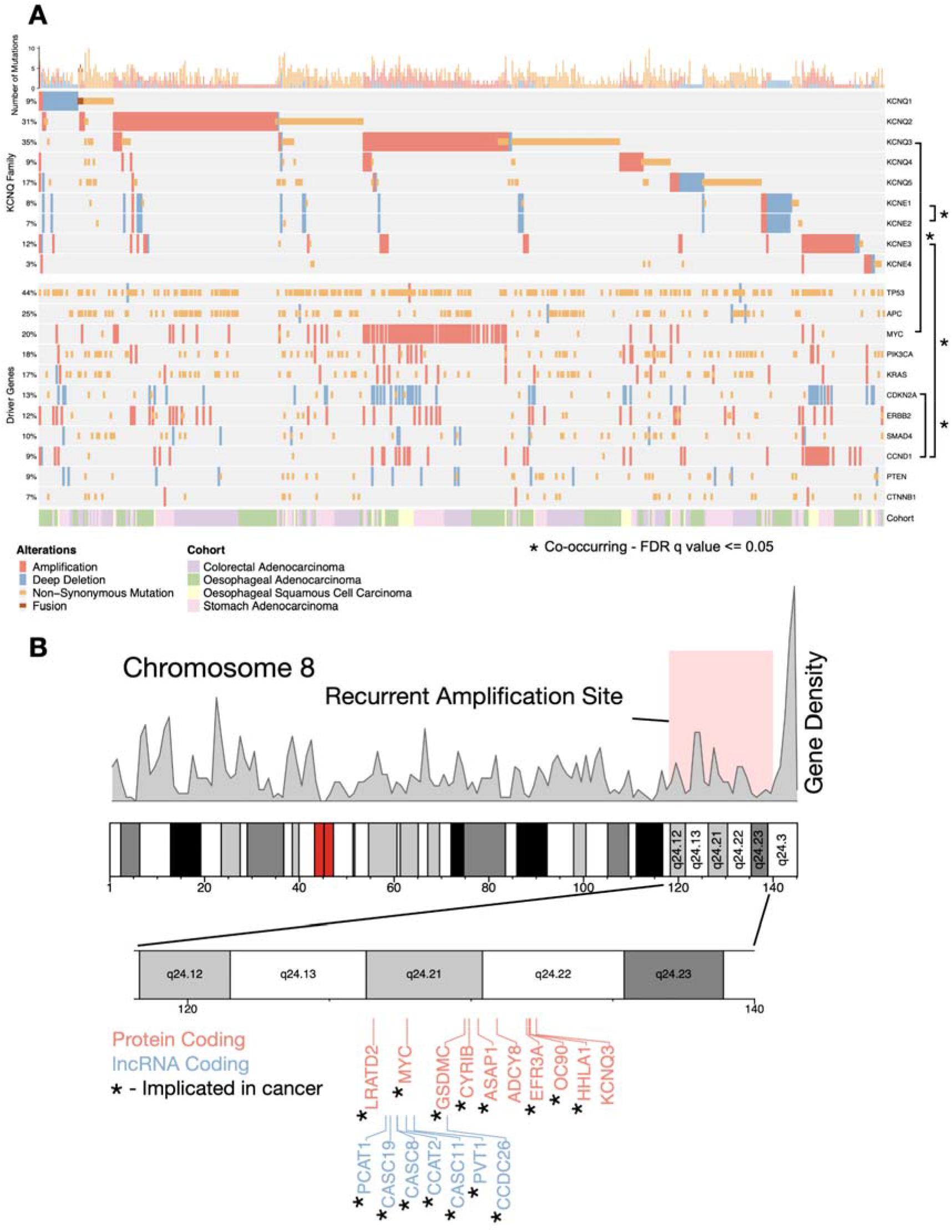
KCNQ genes are highly altered in GI cancers. **A)** Oncoprint of genetic alterations in KCNQ/E gene family, and a set of known GI cancer driver genes. * represents fdr q value < 0.05 co-occurrence of alterations. **B)** Chromosome 8q24.12-23 showing gene density, and identified genes that are recurrently amplified. * represents genes that are known drivers in human cancer.

Overall our analysis shows that KCNQ alterations are frequent, generally mutually exclusive, and *KCNQ2* and *KCNQ3* are located in known susceptibility loci.

Missense mutations in our cohort are also under evolutionary selective pressure, and most notable genes are *KCNQ1*, which is under negative pressure and generally deleted, and *KCNQ3*, which is under positive selective pressure in OAC, generally amplified, and on a known cancer susceptibility locus.

### Mutations in *KCNQ* genes impact channel function

Having studied types of genetic alterations across GI cancers, we next sought to investigate how missense mutations might alter KCNQ function. Whilst metrics such as dN/dS evaluate selection, this is limited to effects that can be understood from the sequence alone. It follows that if mutations are meaningful, they should be interpretable through changes in the protein structure. KCNQ channels contain 6 transmembrane helices (**Figure S2A**). Helices S1, S2, S3, and S4 make up a voltage sensor domain. The S5, pore, and S6 domain contain the gating components of the channel. To study the functional relevance of missense mutations in KCNQ genes, we performed statistical and biophysical analysis using known structural features. To increase the number of variants for statistical and structural analysis, we extracted all mutations in any KCNQ genes from the Catalogue of Somatic Mutations in Cancer (COSMIC)(19), selecting for mutations occurring within patients from untargeted studies or tissues with any cancer along the GI tract.

We first applied statistical techniques to the 1D protein sequence to look a mutational clustering. Non-random mutational clustering (NMC)(20) applied to the location of mutations in protein sequence identifies significant mutational clusters in *KCNQ1* and *KCNQ3* (**Figure 2A, B**), these correlate with a calculated mutational signature-based observed vs expected ratio applied along the protein sequence. For *KCNQ1*, there is a clear hotspot of selected for mutations within the S2-S3 linker region (cluster 1.1), and within the S6 helix (cluster 1.2), despite the whole gene having an overall dN/dS ratio < 1. *KCNQ3* contains a significant mutational hotspot within the S4 voltage sensor helix (cluster 3.1). Interestingly, mutations found in cluster 3.1 in KCNQ3 S4 (R227Q, R230C, and R236C) are known gain-of-function (GoF) gating mutants implicated in autism spectrum disorders(21,22) (**Table S2**), indicating that cancers are selecting for mutations that increase KCNQ3 channel gating activity, and we thus conclude that some mutations in GI cancer patients increase the activity of KCNQ3. Whilst KCNQ1 and KCNQ3 are the primarily clustered genes, there are additional regions of clustering in some other members of the KCNQ family (**Figure S2B-D**).

**Figure 2:**
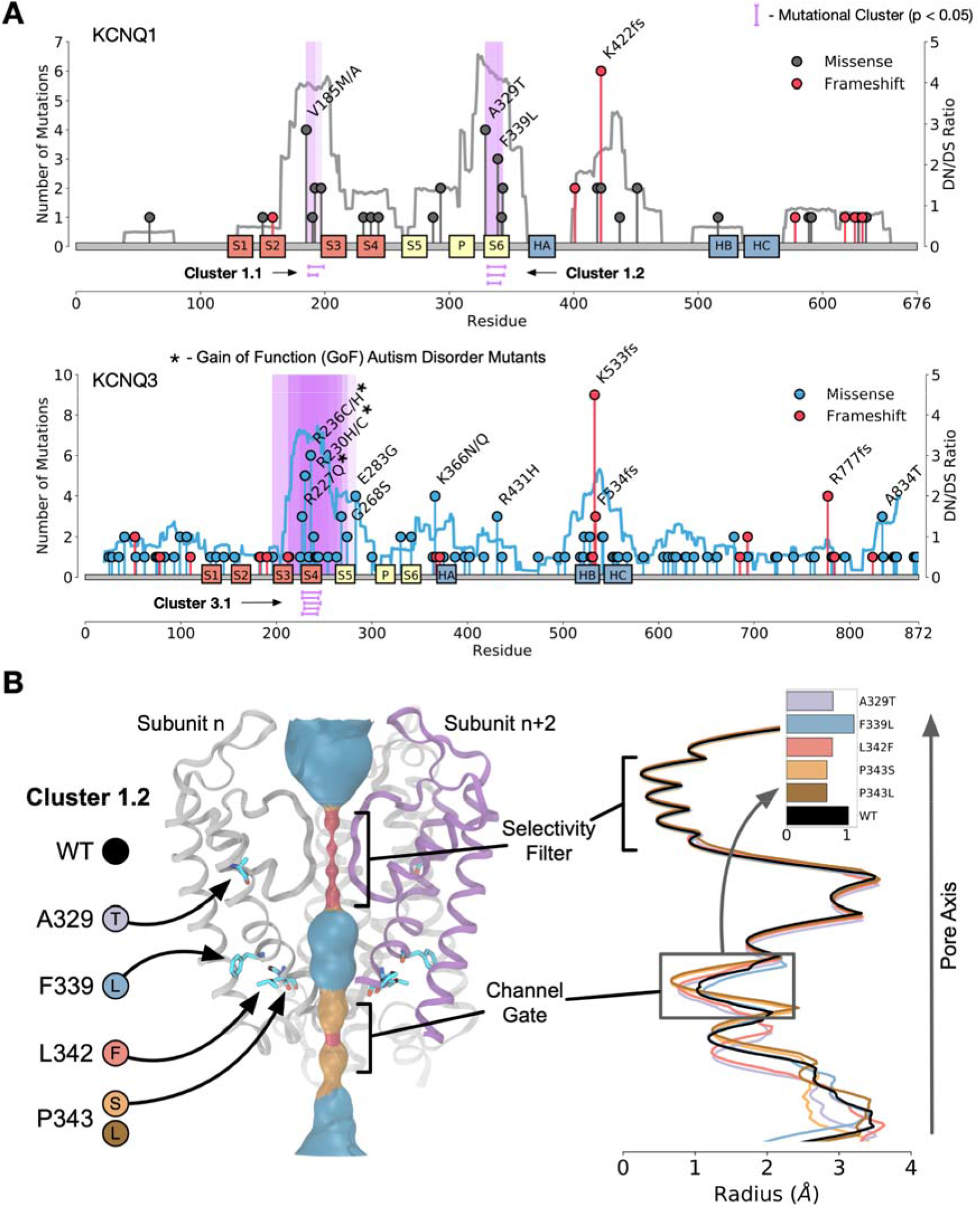
Mutations in KCNQ genes in GI cancers alter channel function. **A)** Mutational clustering for *KCNQ1* (top), *KCNQ3* (bottom), coloured lines represent Observed vs Expected dN/dS ratio, purple highlights represent statistically significant (NMC q-value < 0.05) clusters of mutations. **B)** Render of the pore region of KCNQ1. (left) Mutations modelled are highlighted. (right) HOLE analysis of the pore region of KCNQ1 WT (black) and mutations in cluster 1.2 inset is the smallest distance in the channel gate for each mutation.

To study the structural context of the mutational clusters observed we modelled the atomic 3D structures of KCNQ proteins. Homology models of each human member (KCNQ1-KCNQ5) were generated from the cryo-em structure of *Xenopus laevis* KCNQ1 (protein databank id: 5VMS) and simulated for 200ns using atomistic molecular dynamics in a POPC membrane to validate model soundness (**Figure S2E**). Overlaying mutational frequency with the structures shows areas of high mutational burden, notably the S4 helix of KCNQ3 (**Figure S2F**). Calculation of mutational clusters in the 3D structures of each protein also reveals a statistically unlikely (p <0.05) distribution of two clusters in KCNQ1 (**Figure S2G**), one of which is in the pore region (overlapping with cluster 1.2), and the other of which is in a known phosphatidylinositol (PIP) binding regulatory site(23), the disruption of which would reduce gating activity. As mutations in cluster 1.2 in KCNQ1 are in the vicinity of the pore we generated models for each variant - F339L, L342F, P343L, and P343S, and an additional frequently observed mutant (A329T) (**Figure 2C**) within a single subunit of the channel. Pore diameter calculations show that all mutations except F339L occlude the pore, reducing or eliminating its ability to gate potassium ions, even when a single subunit is mutated, and so we conclude that mutations in cluster 1.2 are likely loss-of-function (LOF).

### KCNQ channels control cell proliferation and correlate with clinical outcome

Based on the apparent links between *KCNQ* genomic status and cancer from patient data, we next sought to establish how changes in *KCNQ1* and *KCNQ3* expression impacts cancer cell phenotype. RNA expression analysis across our cohort (n=1594) finds that *KCNQ1* is generally downregulated in GI cancers vs normal tissue, and *KCNQ3* is significantly upregulated at the RNA level in both oesophageal and colorectal adenocarcinoma (**Figure 3A**) – consistent with the patterns of amplification and deletion observed previously. Cox proportional hazards ratio (**Figure 3B**) also highlights a significant (p < 0.05) positive correlation between patient outcome and *KCNQ1* expression, and a negative correlation (p = 0.07) between outcome and *KCNQ3* expression when tissue differences are included. Looking specifically at OAC, top and bottom quartiles of *KCNQ3* expressors have significantly different survival outcomes (**Figure 3C**), with patients expressing more *KCNQ3* having a worse overall survival.

**Figure 3:**
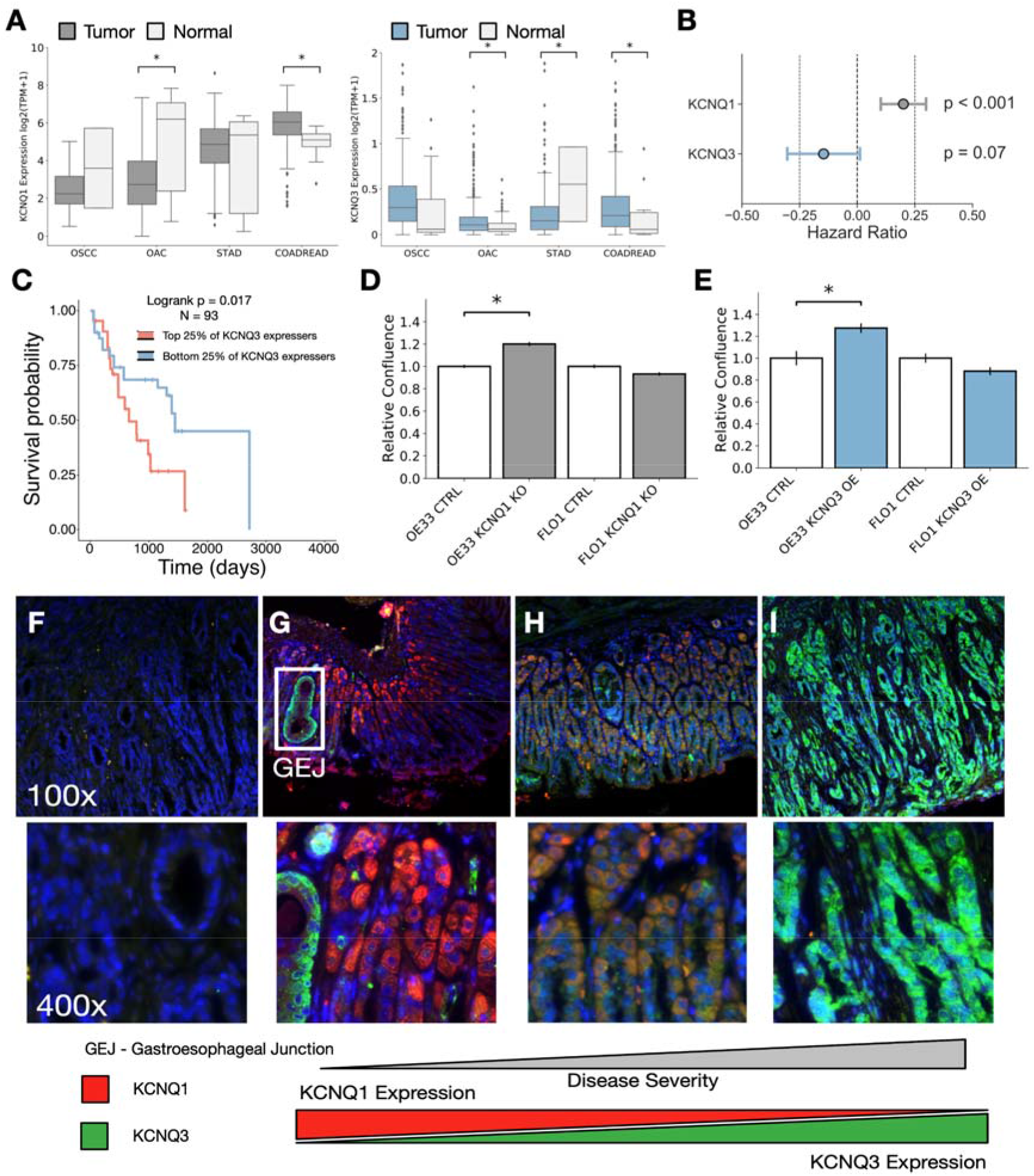
KCNQ expression alters GI cancer cell phenotype. **A)** RNAseq expression for KCNQ1 and KCNQ3 in our patient cohorts. **B)** Multivariate Cox-Regression analysis of KCNQ1 and KCNQ3 in GI cancers. **C)** Kaplan-Meier analysis of upper and lower quartile patients with Oesophageal Adenocarcinoma subset by KCNQ3 gene expression. **D)** Relative confluence of cell growth in WT vs KCNQ3 overexpressing (OE) OE33 and FLO1 cell lines. **E)** Relative confluence of cell growth in WT vs KCNQ1 knockout (KO) OE33 and FLO1 cell lines. **F-I)** images from mouse stomach tissue. Blue represents celltiter blue, red represents KCNQ1, and green represents KCNQ3. Images shown are: **F, G)**, Normal Stomach **H),** Benign Adenoma **I)**, Metastatic Adenocarcinoma.

Due to the co-occurrence of KCNQ3 and MYC amplification it is difficult to distinguish between effects solely caused by increased expression of KCNQ3 rather than an amplification of chr8q24, and so we chose to experimentally evaluate if the expression of KCNQ genes can impact cancer cell phenotype in the most consistently associated cancer subtype – OAC. We chose to reduce expression of KCNQ1 using a CRISPR-Cas9 induced knockout (KO)(**Figure S3A**), and overexpress (OE) KCNQ3 in Oesophageal Adenocarcinoma cell lines OE33 and FLO-1 (**Figure S3B, C**). KO of KCNQ1 significantly increases the proliferation rate (p < 0.05) of OE33 cell lines (**Figure 3D**), but does not change proliferation rate in FLO1 cells. KCNQ3 similarly significantly increases proliferation rate (p < 0.05, though induces a small decrease in cell size – **Figure S3C**) when overexpressed in OE33 (**Figure 3E**), but induces a small decrease in proliferation in FLO1 cells. This suggests KCNQ1 expression can supress OAC proliferation, and KCNQ3 expression can promote it, prompting us to study an *in vivo* model which may be more functionally relevant.

To bolster our findings and explore their generality, we looked to a murine *Prom1^C-L^;Kras^G12D^;Trp53*^*flx*/flx^ model of GI cancer(24). *Prom1* marks a stem compartment of progenitor cells that replenish tissue and cause cancers of the GI tract when mutated. Comparing the transcriptomes of isolated *Prom1*+ gastric stem cells and their *Prom1*- daughter cells from normal gastric mucosa and gastric adenocarcinomas, we observe that *KCNQ1* is downregulated and *KCNQ2/3/5* genes are significantly upregulated (q <0.05) in gastric adenocarcinomas (**Figure S3D**).

To validate these changes we immunostained for KCNQ1 and KCNQ3 in *Prom1^C-L^;Kras^G12D^;Trp53*^*flx*/flx^ murine gastric mucosal tissue. Normal gastric mucosa weakly expresses KCNQ3 (green), and has moderate expression of KCNQ1 (red) (**Figure 3F, G).** In benign adenoma tissue (**Figure 3H**) there is an upregulation of KCNQ3 and slight decrease in KCNQ1. In metastatic adenocarcinoma there is an almost complete loss of KCNQ1, and concurrent upregulation of KCNQ3 (**Figure 3I**), confirming that KCNQ protein levels correlate with disease severity in a model of GI cancer. We additionally find a weak but significant correlation between *KCNQ1* and *KCNQ3* expression and tumor stage in patient data of GI cancer, suggesting that this finding may be extended to human cancer (**Figure S3E**).

### KCNQ activity mediates Wnt, Beta-catenin, and MYC signalling through cadherins junctions

We next looked to understand how the expression of KCNQ genes impacts major cancer signalling pathways. We calculated the PROGENY pathway scores for every patient with RNA expression data, and correlated the scores of all 14 pathways with KCNQ1 and KCNQ3 gene expression through a linear regression model, correcting for tissue specific differences (**Figure 4A, S4A**). We find a significant correlation between KCNQ expression and the interlinked PI3K, Wnt, TGFb, and TNFa pathways. We also confirm an established link between KCNQ1 and beta-catenin signalling in patients, as well as predicting a similar relationship with KCNQ3, as clustering patients based on Wnt pathway genes finds a statistically significant partitioning of patients by high and low KCNQ expression (**Figure S4B**).

**Figure 4:**
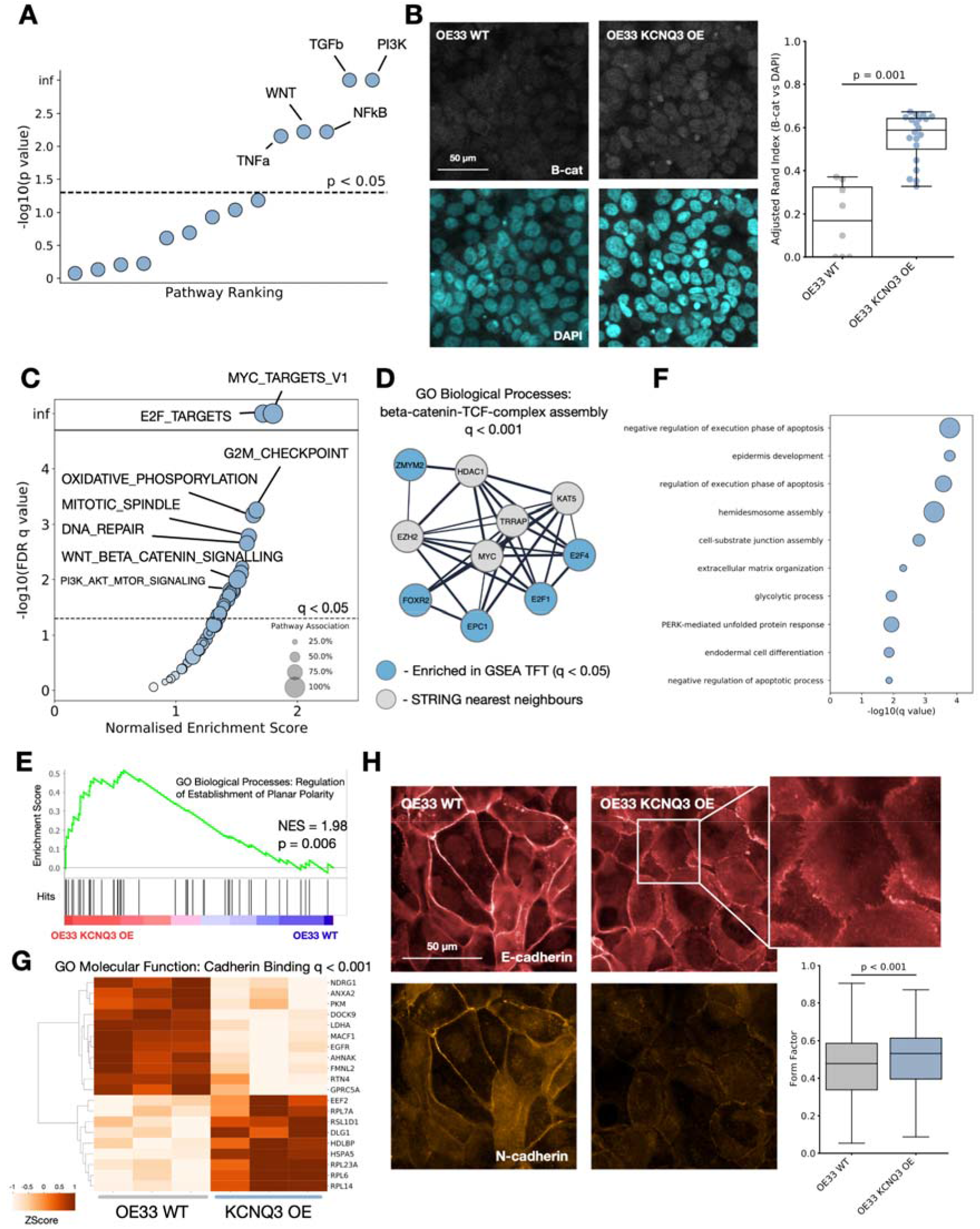
KCNQ activity mediates beta-catenin signalling. **A)** Progeny pathway correlation significance with KCNQ3 RNA expression **B)** Imaging of beta-catenin localisation (top – silver), and nuclear staining (bottom – blue) for WT OE33 (left) and KCNQ3 overexpression (OE) OE33 cell lines, (right) **C)** Enrichment of Hallmarks gene sets by GSEA for WT vs KCNQ3 OE OE33 cells. **D)** String analysis of top 5 transcription factors identified by GSEA TFT gene sets. Genes enriched for GO Biological Processes identify Beta-Catenin signalling q<0.001. **E)** GSEA enrichment plot for GO Biological Processes: Regulation of Establishment of Planar Cell Polarity applied to WT OE33 cell lines vs KCNQ3 OE OE33. **F)** GO Biological Process enrichment significance for significantly (q < 0.05) differentially expressed genes in KCNQ3 WT vs KCNQ3 OE OE33 **G)** Heatmap of genes involved in the most enriched GO molecular function (Cadherin binding) for KCNQ3 WT vs OE OE33. **H)** Imaging of E-cadherin (red) and N-cadherin (orange) in WT OE33 (left) vs KCNQ3 OE OE33 (right), and form factor calculation for microscopy images, N = 1371.

To validate the prediction that KCNQ3 activity may interact with the Wnt pathway, and to deconvolute KCNQ3 expression and MYC amplification in patients, we stained for the localization of Beta-catenin in our KCNQ3 modulated cell lines. OE33 cells overexpressing KCNQ3 show a significantly stronger nuclear localization of Beta-catenin (median Adjusted Rand Index overlap B-cat and DAPI increase of 0.4, p < 0.05) when compared to WT OE33 (**Figure 4B**). FLO1 cells are known to already have a basal Beta-catenin activity(25) which may offer an explanation for why FLO1 KCNQ modified cells do not show significant proliferation increases. To further study the effect of KCNQ in GI cancers, we performed RNA sequencing analysis on our modulated and WT OE33 cell lines. Gene set enrichment analysis (GSEA)(26) confirms a significant positive enrichment for beta-catenin signalling in the KCNQ3 OE and KCNQ1 KO cell lines (**Figure 4C, Table S3**), as well as MYC signalling, E2F transcription factor activity, and G2M checkpoint activity – consistent with a more proliferative phenotype. Transcription factor enrichment against the TFT gene set identifies a series of transcription factors linked to MYC, and that overlap significantly (q < 0.05) with beta-catenin signalling (**Figure 4D, S4C**). Interestingly, as KCNQ3 is recurrently amplified alongside MYC, this suggests that KCNQ3 may act as an amplifier of MYC in this context, similarly to the recently identified lncRNA *PVT1*(27). Finally, GSEA against the GO biological processes set identifies a significant enrichment for planar cell polarity pathways, and non-canonical Wnt signalling (**Figure 4E, Table S4**) – a subtype of Wnt signalling associated with maintenance of cell polarity, and known to play a role in cancer(28).

To further study pathways altered in our cell lines, we performed differential expression analysis followed by enrichment. Enrichment for GO biological processes on differentially expressed (q <0.05) genes (**Table S5**) identifies biological processes including apoptosis control, cellular junctions, and cell development differentiation (**Figure 4F**), and clusters of differentially expressed pathways including MYC and Wnt signalling, NFKB signalling, and protein kinase C (**Figure S4D**). The top enriched GO molecular function in KCNQ3 OE is cadherin binding (**Figure 4G, S4E**), consistent with a mechanism of action where KCNQ activity alters the structure of cadherins junctions and changes the signalling activity of beta-catenin, as well as potentially activating other pathways such as NFKB or planar cell polarity.

To explore how KCNQ3 might influence planar cell polarity we immunostained for the presence of E-cadherin and N-cadherin (CDH1 and CDH2) in our OE33 cell lines (**Figure 4H**), and discovered that KCNQ3 OE results in a change in cadherin expression and cellular morpohology. KCNQ3 OE OE33 are more rounded (median Form Factor difference of 0.05, p < 0.05, N = 1371), and many cells show the presence of membrane ruffles when E-cadherin is stained. Membrane ruffles have been observed previously and are associated with changes to cell motility and extracellular matrix organisation, consistent with our RNAseq analysis. We also find that N-cadherin expression is decreased, showing that this change is more complex than traditional epithelial-to-mesenchymal transition (EMT).

### KCNQ channels have therapeutic potential in GI cancer

Having identified that KCNQ expression induces cancer-associated changes in OE33 cells, we next sought to confirm these findings in patient data. We compared GO biological process terms associated with significantly differentially expressed genes (q < 0.05) for OE33 KCNQ3 OE vs WT, and for the 25 highest and lowest KCNQ3 expressing patients with Oesophageal Adenocarcinoma (**Table S6,7**). We find a significant (p < 0.0001) overlap between pathways altered in our cell lines vs patients (**Figure 5A**). Moreover, there is almost complete overlap (98.5%) between pathways altered in OE33 KCNQ3 OE cell lines and patients, indicating that our cell lines accurately reproduce a subset of patient relevant, cell autonomous pathways. Ranking pathways by average q-value, (**Figure 5B**) the top 10 pathways include differentiation and development pathways, extracellular matrix organisation, and cell migration pathways, suggesting that patients overexpressing KCNQ3 result in similar disruptions to cellular development and morphology as in OE33 KCNQ3 OE cells. We find a similar trend is observed when KCNQ1 KO vs WT pathways are compared with the top and bottom 25 OAC patients by KCNQ1 expression (**Figure S5A, B, Table S8,9**), (overlap between cell lines and patients of 87.1% and 65.9% respectively, overlap p < 0.001) suggesting that patients with low *KCNQ1* expression alter similar pathways to KCNQ1 KO OE33.

**Figure 5:**
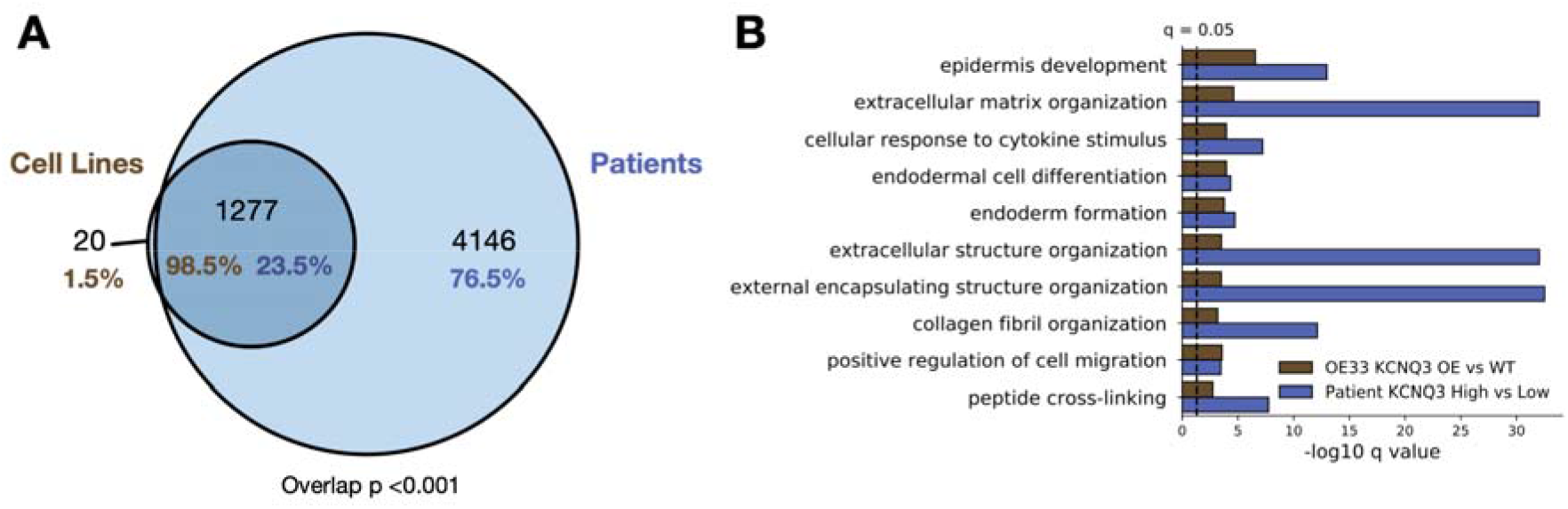
Patients high and low for KCNQ3 alter similar signalling pathways to OE of KCNQ3 in OE33. **A)** Venn diagram of overlap between enriched pathways in cell lines (KCNQ3 WT vs KCNQ3 OE OE33), and patients (highest 25 vs lowest 25 patients by KCNQ3 expression in OAC). Overlap p represents using cell line pathways as custom set in g-proflier. **B)** -log10 q values for the top 10 overlapping pathways between cell lines and patients.

We surmised that modulation of KCNQ1 and 3 activity with small molecules may also confer a therapeutic benefit in GI cancers. KCNQ3 represents a more feasible clinical target than KCNQ1, as it is not involved in the cardiac action potential, and a number of KCNQ3 inhibiting drugs already exist. We applied two drugs to KCNQ3 OE and WT OE33 cell lines, the KCNQ2/3 specific inhibitor linopirdine(29), and the more broad inhibitor amitriptyline(30), which inhibits a large number of proteins and is commonly used to treat depression(31).

Proliferation of both WT and KCNQ3 OE OE33 was significantly reduced upon exposure to linopirdine (**Figure 6A**), and this effect is sensitised by overexpression of KCNQ3. For FLO1 cells however, which did not respond to KCNQ3 overexpression, linopirdine does not have any effect – adding weight to the effect of the drug on cellular proliferation in OE33 being likely due to its actions on KCNQ3. We also find application of amitriptyline has a potent inhibitory effect on growth in OE33 cells, but that this effect is also present in FLO1. This suggests that amitriptyline likely also acts through mechanisms other than KCNQ3 to reduce growth rate. To confirm that linopirdine and amitriptyline mechanism of action involves inhibition of KCNQ3 we also performed RNA sequencing on OE33 cells exposed to 100mg/ml of each drug. There is a strong overlap in the differentially expressed genes associated with application of each drug, with the KCNQ2/3 specific inhibitor linopirdine altering a subset (64.1%) of genes altered by the more broadly inhibiting amitriptyline (**Figure 6B, Table S10,11**). Pathway enrichment and clustering with REVIGO for the overlapping gene sets(32) (**Figure 6C, Table S12**) identifies pathways involved in the cell cycle, WNT signalling, NFKB signalling, and the cytoskeleton as being altered in response to application of either drug, confirming that application of these inhibitors impacts cancer cell phenotype through our proposed mechanisms. We also find cadherins junctions are amongst the most enriched GO molecular functions in both instances (**Figure S6A, B**). Finally, to confirm a reduction in MYC/WNT signalling in OE33 exposed to drugs, differential expression identifies significant (q < 0.05) reduction in the expression of downstream responders MYC, Cyclin D1, E-cadherin, and E2F1, (**Figure 6C**) all of which are known players in the progression of GI cancer.

**Figure 6:**
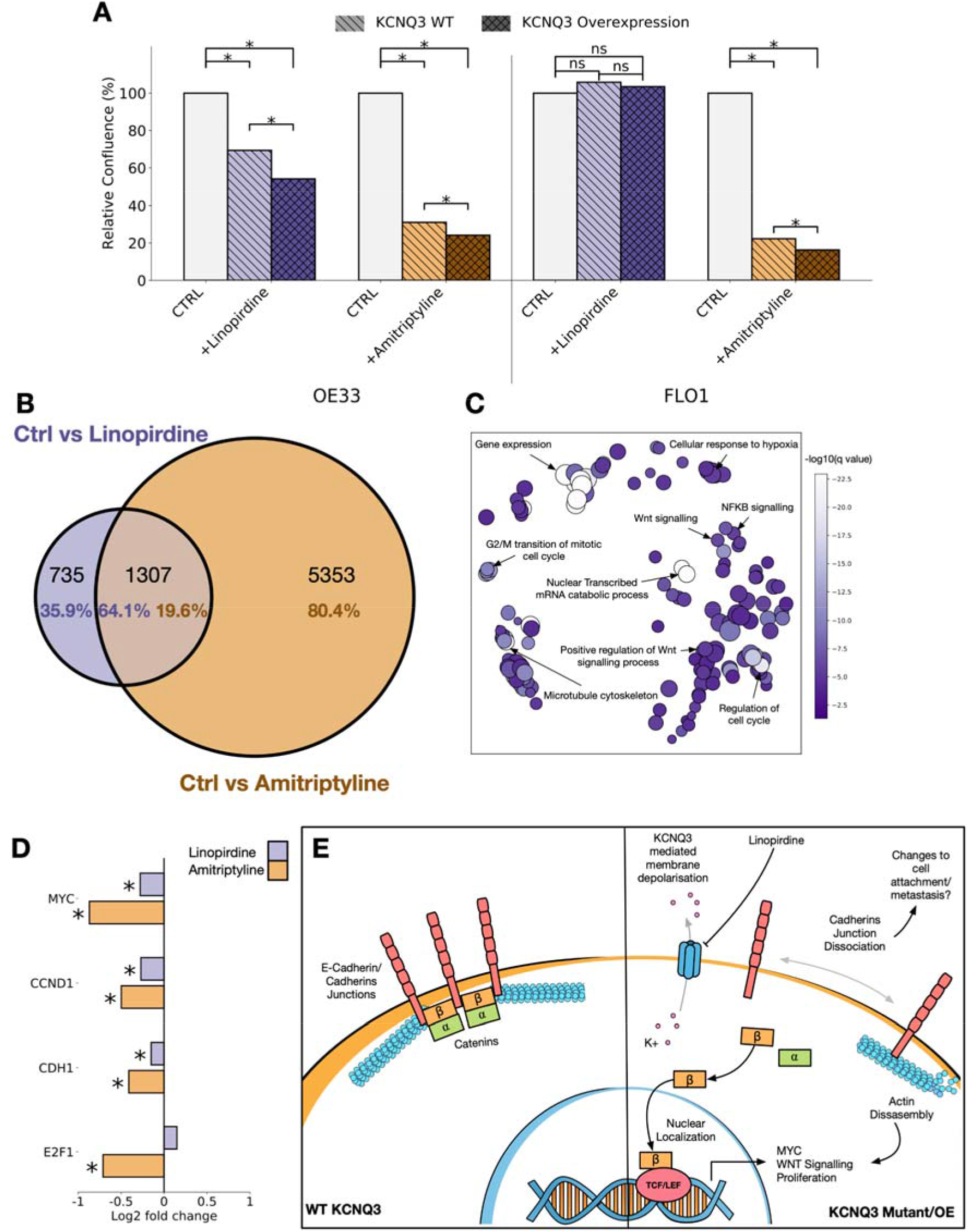
KCNQ channels are potential therapeutic targets in GI cancer. Relative confluence plots for OE33 (left) and FLO1 (right) cell lines exposed to linopirdine (purple) or amitriptyline (orange). Cell lines are either WT for KCNQ3 (light), or KCNQ3 overexpressing (OE) (dark). **B)** Venn diagram of overlapping GO biological processes enriched in ctrl vs linopirdine exposed KCNQ3 OE OE33 cells (purple) and ctrl vs amitriptyline exposed KCNQ3 OE OE33 cells. **C)** REVIGO clustered GO biological process terms associated with overlap between ctrl vs linopirdine and amitriptyline exposed KCNQ3 OE OE33 cells. **D)** Fold change of MYC, CCND1, CDH1, and E2F1 in ctrl vs linopirdine exposed (purple) and ctrl vs amitriptyline exposed (orange) KCNQ3 OE OE33 cells. * represents q value < 0.05. **E)** Suggested mechanism of KCNQ3 activity on GI cancer cells, and their inhibition by linopirdine.

## DISCUSSION

There is emerging evidence that ion channels play a role in many, and potentially all cancers(33). Therapeutics against voltage gated potassium channels improve prognosis for glioblastoma and breast cancer(34), and studies implicate specific sodium(35) and calcium channels(36) in cancers. We show that the *KCNQ* family of genes play a significant and functional role in human gastrointestinal cancers. Through integration of data at the patient, cell, and protein structural levels, coupled with *in vitro* models we show that *KCNQ* genes and protein products contribute to cancer phenotype and are a potential therapeutic target. We show that a large number of patients with gastrointestinal cancers have genetic alterations in a member of the *KCNQ* family, and expression levels of these genes are associated with patient outcome. Mutations in the *KCNQ* family have functional effects on the protein and are under selective pressure, and we find that *KCNQ* activity controls signalling activity of the WNT pathway through changing the localization of beta-catenin, and drives the cell cycle and MYC activity, as well as having a role in cell polarity. We propose a mechanism for KCNQ3 activity in GI cancer, whereby it controls the clustering/assembly of cadherins junctions. Dissociation of these junctions through changes in the membrane potential controlled by KCNQ3 may lead to the activation of WNT and MYC signalling, as well as changing cellular polarity and morphology (**Figure 6D**). Finally, we demonstrate that *KCNQ* family members are a viable drug target with the use of already available therapeutic compounds that have not yet been actioned against cancer, but have been FDA approved for other uses. This is particularly interesting in the case of KCNQ3 – as it is often recurrently amplified with MYC and can independently drive MYC activation, it may act as a gateway to modulating the notoriously hard to drug MYC signalling in patients(37,38), but we caution that correlations of KCNQ with patient survival may be heavily biased by MYC convolution, amidst other emerging problems identified with survival analysis(39).

By studying data from varying sources simultaneously we find consistently that *KCNQ1* exhibits properties of a tumor suppressor – it is often deleted or lost in patients, mutations are generally inactivating, and cell proliferation can be increased when it is lost. Opposingly, *KCNQ3* shows hallmarks of an oncogene, it is often amplified in cancers, mutations are mostly GoF, and cell proliferation/wnt signalling increases when it is overexpressed. Thus genes within the same family, with very similar molecular activity have apparently opposing influences on cancer phenotype. Caution must thus be taken when considering therapeutic applications of KCNQ involvement in cancer, due also to the extreme importance of KCNQ1 in cardiac activity. Despite this, existing compounds specific to KCNQ3(40) may have a therapeutic window, as is thought to be the case with hERG inhibitors(41). That the KCNQ2/3 specific inhibitor linopirdine shows no effect in FLO1 cell lines, but they are potently inhibited by the broader acting amitriptyline indicates a key role for a number of other proteins that may be therapeutic targets in GI cancer, but also opens up the possibility that patients already taking amitriptyline as an analgesic/antidepressant in cancer may be impacted by its other effects.

## METHODS

### Data acquisition

TCGA level 3 data was downloaded using Firebrowse (RNAseq) or cBioportal (copy number alteration, mutation and clinical data)(42). COSMIC data(19) was downloaded from cancer.sanger.ac.uk (version 92). We subset mutations into those only found in gastrointestinal tissue, defined as those where the primary site is in one of the following categories: “large_intestine”, “small_intestine”, “gastrointestinal_tract_(site_indeterminate)”, “oesophagus”, “stomach”.

### Oncoprint

Oncoprint was generated using the oncoprint library in R(43). Copy number alterations were determined as follows – relative copy number for each gene was defined as:

> 〖relative copy number= log〖 _2 ((Total Copy Number of Gene)/(Total ploidy of sample))

Genes were defined as deleted if total copy number == 0 OR relative copy number < −1, genes were defined as amplified if relative copy number > 1.

### Co-association analysis

Co-association analysis was performed using DISCOVER(44).

### Chromosome plots

Chromosome plots were generated using the karyoplotR library(45).

### Dn/Ds

Dn/Ds for individual genes was calculated using the dndscv library applied to all mutations across all cancers grouped togther, as well as to each tissue (OSCC, OAC, STAD, COADREAD)(46).

For calculating the expected vs observed mutational distribution, exon data was downloaded from the Ensembl Biomart (ensembl.org/biomart). Ensembl 96, hg38.p12 was selected, and data downloaded for chromosomes 1-22, X, and Y. We used bedtools(47) to sort data and overlapping exons were merged. To sort data we used the following command:

~~~
tail -n +2 human_exon_bed_1-Y.txt | cut -f 1,2,3 | bedtools sort -i stdin > human_exon_bed_file_l-Y_sorted.bed
~~~

Merging was performed using:

~~~
bedtools merge -i human_exon_bed_file_1-Y_sorted.bed > human_exon_bed_file_1-Y_merged.bed
~~~

The R library Deconstructsigs(48) was used to generate the mutational signature for all COSMIC mutations in KCNQ genes within GI tissue for these exons. The mutational spectrum was normalised using the mutational signature to generate the expected relative mutation rate for each possible missense mutation. This was then multiplied by the total number of mutations in each gene to get the expected distribution of events along the gene. The observed and expected mutational frequency ratio was average over a sliding window of 50 bases.

### NMC

Mutational clustering was calculated using the NMC clustering method from the R library IPAC(20) applied to sequence alone. All mutations to each KCNQ gene in the COSMIC database for GI cancers were considered. The top 5 mutational clusters ranked by adjusted P-value were plotted.

### Cox proportional Hazards/KM

Cox proportional hazards was performed using the python library lifelines (49). Patients were labelled with their cancer origin (OAC, OSCC, STAD, COADREAD), and overall survival was correlated with zscored rna expression of KCNQ1, KCNQ3, and the previously studied driver genes (APC, MYC, TP53, SMAD4, PIK3CA, KRAS, CDKN2A, CTNNB1, ERBB2, CCND1, PTEN) concurrently.

Kaplan Meier analysis was performed using the R library Survival(50). We used the clinical data associated with the TCGA and OCCAMS, which includes overall survival. Survival times were converted to days, survival curves were generated on the top and bottom 25% of patients ranked for expression of KCNQ3.

### Homology Modelling/MD simulations

Homology modelling was performed using the template structure 5VMS from the protein data bank(51). Sequences were aligned with MUSCLE (52) before manual adjustment based on key residues (arginines in the S4 helix, key regions of the pore domain. Single point mutations were induced in the models using the mutate_model protocol in the modeller tool as described in Feyfant et al (53), and using the mutate_model.py script available on the modeller website.

Molecular dynamics was performed using Gromacs version 2018.1 (54).

For simulations of homology models in AT we used the charmm36 forcefield (55). In each case the protein was placed in a 15 x 15 x 15nm box with roughly 650 DPPC lipid molecules. Setup was performed in the same manner as systems in the memprotMD pipeline (56). The system was converted to MARTINI coarse-grained structures (CG-MD) with an elastic network in the martiniv2 forcefield(57) and self-assembled by running a 1000ns molecular dynamics simulation at 323k to allow the formation of the bilayer around the protein. The final frame of the CG-MD simulation was converted back to atomistic detail using the CG2AT method (58). The AT system was neutralised with counterions, and additional ions added up to a total NACL concentration of 0.05 mol/litre. The system was minimized using the steepest descents algorithm until maximum force Fmax of the system converged.

Equilibration was performed using NVT followed by NPT ensembles for 100 ps each with the protein backbone restrained. We used the Verlet cutoff scheme with PME electrostatics, and treated the box as periodic in the X, Y, and Z planes. Simulations were run for 200ns of unrestrained molecular dynamics. Root mean square deviation (RMSD) was calculated for structures using the g_rmsdist command in GROMACS.

CG simulations of single helices were performed as described previously (59). Models of single helices were generated and converted to MARTINI coarse grained structures. Helices were then inserted into POPC bilayers and simulated for short (100ns) simulations for 100 repeats of each sequence.

### Pore calculations

Pore analysis was performed using the algorithm HOLE (60). Pore profile was visualised using Visual Molecular Dynamics (VMD) (61).

### RNAseq processing

Quality control of raw sequencing reads was performed using FastQC v0.11.7. Reads were aligned to GRCh37 using STAR v2.6.1d. Read counts were generated within R 3.6.1 summarizeOverlaps.

### GSEA

Gene set enrichment was performed using the GSEA desktop application (62). GSEA was run for 5000 permutations, and phenotype permutations were used where the number of samples was lower than 7, otherwise gene set permutations were performed.

### Differential expression

Differential expression analysis was performed using the R library Deseq2 (63), performed on count data. All analysis was run to compare two groups, groups were assigned within a condition matrix, and the analysis run using the formula:

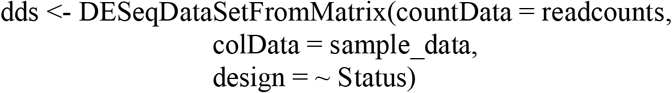

### Cell culture

OE33 was cultured in RPMI (Roswell Park Memorial Institute) 1640 medium supplied with 10% FBS (fetal bovine serum) and FLO1 was cultured in DMEM (Dulbecco’s Modified Eagle Medium) supplied with 10% FBS.

### CRISPR knockout of KCNQ1

We first generated CRSRP-Cas9 expressing cell lines of OE33 and FLO1. Briefly, Lentiviral particles were generated by transfecting HEK293T cells with a Cas9 expressing plasmid, FUCas9Cherry, gift from Marco Herold (Addgene plasmid # 70182 (64)), an envelop plasmid, pMD2.G and a packaging plasmid, psPAX2, both gifts from Didier Trono (Addgene plasmid # 12259 and #12260). OE33 and FLO1 cells were transduced with the lentivirus, sub-cultured and selected for mCherry^bright^ cells using FACS, respectively, to generate stable Cas9 expressing cell lines.

We then designed four sgRNA sequences that targeting exon2 and exon3 of KCNQ1, which were shared by both known KCNQ1 variants, using an online tool http://crispor.tefor.net/. Namely, sequences #6 CAGGGCGGCATACTGCTCGA and #7 GGCGGCATACTGCTCGATGG targeting exon2; and #8 GGCTGCCGCAGCAAGTACGT and #9 CGGCTGCCGCAGCAAGTACG targeting exon3. sgRNA sequences were cloned into a backbone plasmid pKLV2-U6gRNA5(BbsI)-PGKpuro2ABFP-W (Fig S3A, left panel), which was a gift from Kosuke Yusa (Addgene plasmid # 67974), as described in [PMID: 27760321]. Briefly, for each sgRNA, two complimentary oligos were purchased, annealed and cloned into the BbsI site of the backbone plasmid pKLV2. sgRNA lentivirus were then generated using the aforementioned pMD2.G and psPAX2 plasmids from HEK293T cells.

OE33 and FLO1 Cas9 expressing cell lines were transduced with the four sgRNA lentivirus to generate KCNQ1 knockout cell lines, which were later sub-cultured and selected by puromycin treatment of 5 μg/mL for 3 days. Four KCNQ1 knockout cell lines were generated using the four sgRNA sequences. Genomic DNA were extracted from each cell lines using Qiagen AllPrep DNA/RNA Kit, the sgRNA targeted regions were amplified in PCR using ACCUZYME™ DNA polymerase (BIO-21052, Meridian Bioscience) according to the manufacturer’s manual. Primers for the PCR were: TCCCCAGGTGCATCTGTGG (forward) and TCCAAGGCAGCCATGACAT (reverse) for sgRNA sequences #6 and #7 targeting exon2; and TGCAGTGAGCGTCCCACTC (forward) and CTTCCTGGTCTGGAAACCTGG (reverse) for sgRNA sequences #8 and #9 targeting exon3. PCR products were approximately 200 bp long, which were run in 1% agarose gel and purified using Qiagen Gel Extraction Kit, and then sent for Sanger Sequencing provided by Source BioScience. Successful KCNQ1 knockout by the non-homologous DNA end joining (NHEJ) was confirmed in the cell line used sgRNA #9 (Fig S3A, right panel).

### Overexpression of KCNQ3

To generate KCNQ3 overexpressing lentiviral plasmid, KCNQ3 fragment was cloned from pCMV6-KCNQ3 (RC218739, OriGene) using ACCUZYME™ Mix 2x (BIO-25028, Meridian Bioscience) using primers of GGGCCTTCTAGAATGAAGCCTGCAGAACACGC (forward, with a XbaI cloning site) and TCACACGCTAGCTTAAATGGGCTTATTGGAAG (reverse, with a NheI cloning site).

The KCNQ3 PCR product was purified using Qiagen QIAquick PCR Purification Kit, and then cloned into a backbone plasmid with a EGFP tag, pUltra, a gift from Malcolm Moore (Addgene plasmid # 24129, (65)) between the XbaI and NheI sites (Fig. S3B). Lentivirus were then generated using the aforementioned pMD2.G and psPAX2 plasmids from HEK293T cells, which were used to transduce OE33 and FLO1 cells. Stable KCNQ3 overexpression cell lines were then generated from sorting for EGFP^birght^ cells using FACS.

### Proliferation assay

Cells of knockout or overexpression were seeded in 24 well plate at 50,000 cells per well. 4 replicates per cell type or drug treatment condition. Plates were cultured in IncyCyte SX5 and scanned for the whole well using the Standard Phase model every 6 hours. Cell confluence were quantified using the built-in Basic Analyzer and plotted over time. Each experiment was repeated for at least once.

### Western blot

Cells were freshly harvested and counted. 600,000 cells were lysed using lysis buffer containing 50% of TruPAGE™ LDS Sample Buffer (PCG3009, Merck) and 20% 2-mercaptoethanol. Cell lysates were heated at 98C for 5 minutes, cooled down to room temperature, diluted 1:1 using water and run in NuPAGE™ precast gels (NP0321BOX, ThermoFisher Scientific). The gels were transferred to membranes using iBlot system (IB401001, ThermoFisher Scientific). The membranes were incubated with primary antibodies KCNQ3 (GTX54782, GeneTex, 1:1000) and GAPDH (ab181602, Abcam, 1:10,000) at 4C overnight, followed by IRDye® 800CW Goat anti-Rabbit IgG Secondary Antibody (925-32211, Li-Cor, 1:5000). The membranes were visualized using Li-Cor Odyssey® CLx system.

### Drug treatment

Linopirdine (L134-10MG, MW 391.46, Sigma) was prepared in absolute ethanol for 100 mM stock as described in (66). A final concentration of 50 μM was used to treat cells. Amitriptyline hydrochloride (A8404-10G, MW 313.86, Sigma), was prepared in absolute ethanol for 60 mM stock as described in (67). A final concentration of 30 μM was used to treat cells. Culture medium was refreshed every 3 days.

### RNA-seq

Total RNA was extracted from fresh cells or mouse tissues using Qiagen RNeasy Mini Kit. RNA-seq libraries were prepared using Lexogen CORALL mRNA-seq kit (098.96 and 157.96) according to the manufacture’s protocol. 3 μg and 700 ng of total RNA input were used for cell line and mouse tissue RNA-seq, respectively. The libraries were sequenced in Illumina NovaSeq platform using SR100. For cell lines, three replicates were sequenced for each cell type or drug treatment condition.

### Murine tissue immunoflouresence

Normal stomach and gastric tumour samples were harvested from *Prom1^C-L^;Kras^G12D^;Trp53*^*flx*/flx^ animals(24). Tissue samples were formalin fixed, paraffin embedded and cut into 5μm sections. Immunofluorescence was performed using sections of formalin fixed, paraffin embedded tissue generated as described above. Antigen retrieval in tissue sections was achieved using pressure-cooking in citrate buffer pH6 for 20 minutes. Tissue sections were incubated with primary antibodies overnight at 4C in a humidity chamber. Primary antibodies included: Kcnq1 (1:50; Abcam, ab77701) and Kcnq3 (1:50; Abcam, ab16228). Following washing, tissue sections where then incubated for 1 hour at room temperature in secondary antibody. Secondary antibodies included Alexa 488 or 594 (1:200; Invitrogen, A-21206 or A-21207). Dual labelling of Kcnq1 and Kcnq3 was performed as sequential stains to account for same species with appropriate single stain controls to monitor for non-specific staining of each antibody. Sections were then counterstained using DAPI (1:10,000; Cell Signaling, 4083) and mounted using ProLong Gold antifade mountant (Thermo Fisher Scientific, P36930). Digital images of tissue sections were captured using a Zeiss ImagerM2 and Apotome microscope.

### Cell line immunofluorescence

Cells were cultured in 8 well chamber slide (154534, ThermoFisher Scientific) to confluence, 4 wells per cell type. The cells were then fixed using 4% paraformaldehyde for 20 minutes at room temperature and blocked using 1% BSA. Immunofluorescent staining was performed using primary antibody of beta-catenin (ab19381, Abcam, 1:100) for overnight at 4C, and secondary antibody of anti-Mouse Alexa Fluor 647 (A-21240, ThermoFisher Scientific, 1:400) for 1 hour at room temperature. Nuclei were counterstained using DAPI. The cells were then imaged using Zeiss LSM 880 confocal microscope with 20x objective.

### Microscopy quantification

Microscopy quantification was performed using cellprofiler3(68). For nuclear vs cytoplasmic beta catenin staining, nuclei were detected using the detect primaryobject command, and their overlap measured using measureimageoverlap tool. For quantification of the cell shape, E-cadherin was used to measure cell shape using the detect primary object command

## ACKNOWLEDGEMENTS

This work was supported by the Medical Research Council (grant no. MR/S000216/1) and through a Grant-in-Aid to the MRC Cancer unit. M.W.J.H. acknowledges support from the Harrison Watson Fund at Clare College, Cambridge. E.R. was supported by Marie Curie, and R.J.G. and E.R. are supported by Cancer Research UK. B.A.H. acknowledges support from the Royal Society (grant no. UF130039).

## REFERENCES

1. Abbott GW. Biology of the KCNQ1 Potassium Channel. New Journal of Science. 2014;

2. Robbins J. KCNQ potassium channels: Physiology, pathophysiology, and pharmacology. Pharmacology and Therapeutics. 2001;

3. Wang JJ, Li Y. KCNQ potassium channels in sensory system and neural circuits. Acta Pharmacologica Sinica. 2016.

4. Ohya S, Asakura K, Muraki K, Watanabe M, Imaizumi Y. Molecular and functional characterization of ERG, KCNQ, and KCNE subtypes in rat stomach smooth muscle. American Journal of Physiology-Gastrointestinal and Liver Physiology. 2015;

5. Millichap JJ, Park KL, Tsuchida T, Ben-Zeev B, Carmant L, Flamini R, et al. *KCNQ2* encephalopathy. Neurology Genetics. American Academy of Neurology Journals; 2016;2:e96.

6. Morita H, Wu J, Zipes DP. The QT syndromes: long and short. The Lancet. 2008. page 750–63.

7. Sands TT, Miceli F, Lesca G, Beck AE, Sadleir LG, Arrington DK, et al. Autism and developmental disability caused by *KCNQ3* gain-of-function variants. Annals of Neurology. 2019;86:181–92.

8. Rapetti-Mauss R, Bustos V, Thomas W, McBryan J, Harvey H, Lajczak N, et al. Bidirectional KCNQ1:β-catenin interaction drives colorectal cancer cell differentiation. Proceedings of the National Academy of Sciences [Internet]. 2017 [cited 2019 Apr 8];114:4159–64. Available from: http://www.ncbi.nlm.nih.gov/pubmed/28373572

9. Fan H, Zhang M, Liu W. Hypermethylated KCNQ1 acts as a tumor suppressor in hepatocellular carcinoma. Biochemical and Biophysical Research Communications. Academic Press; 2018;503:3100–7.

10. Frankell AM, Jammula S, Li X, Contino G, Killcoyne S, Abbas S, et al. The landscape of selection in 551 esophageal adenocarcinomas defines genomic biomarkers for the clinic. Nature Genetics [Internet]. Nature Publishing Group; 2019 [cited 2019 Apr 8];51:506–16. Available from: http://www.nature.com/articles/s41588-018-0331-5

11. Shorthouse D, Riedel A, Kerr E, Pedro L, Bihary D, Samarajiwa S, et al. Exploring the role of stromal osmoregulation in cancer and disease using executable modelling. Nature Communications [Internet]. Nature Publishing Group; 2018 [cited 2019 Apr 8];9:3011. Available from: http://www.nature.com/articles/s41467-018-05414-y

12. Kim J, Bowlby R, Mungall AJ, Robertson AG, Odze RD, Cherniack AD, et al. Integrated genomic characterization of oesophageal carcinoma. Nature. Nature Publishing Group; 2017;541:169–74.

13. Bass AJ, Thorsson V, Shmulevich I, Reynolds SM, Miller M, Bernard B, et al. Comprehensive molecular characterization of gastric adenocarcinoma. Nature [Internet]. Nature Publishing Group; 2014 [cited 2019 Apr 8];513:202–9. Available from: http://www.nature.com/doifinder/10.1038/nature13480

14. Muzny DM, Bainbridge MN, Chang K, Dinh HH, Drummond JA, Fowler G, et al. Comprehensive molecular characterization of human colon and rectal cancer. Nature [Internet]. Nature Publishing Group; 2012 [cited 2019 Apr 8];487:330–7. Available from: http://www.nature.com/articles/nature11252

15. Grisanzio C, Freedman ML. Chromosome 8q24-Associated Cancers and MYC. Genes & Cancer. 2010;1:555–9.

16. Bossé Y, Amos CI. A Decade of GWAS Results in Lung Cancer. Cancer Epidemiology Biomarkers & Prevention. 2018;27:363–79.

17. Integrated genomic characterization of oesophageal carcinoma. Nature. 2017;541:169–75.

18. Martincorena I, Raine KM, Gerstung M, Dawson KJ, Haase K, Van Loo P, et al. Universal Patterns of Selection in Cancer and Somatic Tissues. Cell. Elsevier; 2017;171:1029–1041.e21.

19. Tate JG, Bamford S, Jubb HC, Sondka Z, Beare DM, Bindal N, et al. COSMIC: The Catalogue Of Somatic Mutations In Cancer. Nucleic Acids Research. 2019;

20. Ye J, Pavlicek A, Lunney EA, Rejto PA, Teng CH. Statistical method on nonrandom clustering with application to somatic mutations in cancer. BMC Bioinformatics. 2010;

21. Nappi P, Miceli F, Soldovieri MV, Ambrosino P, Barrese V, Taglialatela M. Epileptic channelopathies caused by neuronal Kv7 (KCNQ) channel dysfunction. Pflügers Archiv - European Journal of Physiology. 2020;472:881–98.

22. Gamal El-Din TM, Lantin T, Tschumi CW, Juarez B, Quinlan M, Hayano JH, et al. Autism-associated mutations in K _V_ 7 channels induce gating pore current. Proceedings of the National Academy of Sciences. 2021;118.

23. Zaydman MA, Cui J. PIP2 regulation of KCNQ channels: biophysical and molecular mechanisms for lipid modulation of voltage-dependent gating. Front Physiol. Frontiers Media SA; 2014;5:195.

24. Zhu L, Finkelstein D, Gao C, Ellison D, Onar-Thomas A, Gilbertson RJ, et al. Multi-organ Mapping of Cancer Risk The generative capacity of an organ’s stem cells determines the life-long risk for developing cancer in that organ. Multi-organ Mapping of Cancer Risk. Cell. 2016;

25. Vangamudi B, Zhu S, Soutto M, Belkhiri A, El-Rifai W. Regulation of β-catenin by t-DARPP in upper gastrointestinal cancer cells. Molecular Cancer. 2011;10:32.

26. Subramanian A, Tamayo P, Mootha VK, Mukherjee S, Ebert BL, Gillette MA, et al. Gene set enrichment analysis: A knowledge-based approach for interpreting genome-wide expression profiles. Proc Natl Acad Sci U S A. 2005;

27. Tseng Y-Y, Bagchi A. The PVT1-MYC duet in cancer. Molecular & Cellular Oncology. 2015;2:e974467.

28. Daulat AM, Borg J-P. Wnt/Planar Cell Polarity Signaling: New Opportunities for Cancer Treatment. Trends in Cancer. 2017;3:113–25.

29. Schnee ME, Brown BS. Selectivity of linopirdine (DuP 996), a neurotransmitter release enhancer, in blocking voltage-dependent and calcium-activated potassium currents in hippocampal neurons. J Pharmacol Exp Ther. 1998;286:709–17.

30. Punke MA, Friederich P. Amitriptyline Is a Potent Blocker of Human Kv1.1 and Kv7.2/7.3 Channels. Anesthesia & Analgesia. 2007;104:1256–64.

31. Moore RA, Derry S, Aldington D, Cole P, Wiffen PJ. Amitriptyline for neuropathic pain in adults. Cochrane Database of Systematic Reviews. 2015;

32. Supek F, Bošnjak M, Škunca N, Šmuc T. REVIGO Summarizes and Visualizes Long Lists of Gene Ontology Terms. PLoS ONE. 2011;6:e21800.

33. Jiang L-H, Adinolfi E, Roger S. Editorial: Ion Channel Signalling in Cancer: From Molecular Mechanisms to Therapeutics. Frontiers in Pharmacology. 2021;12.

34. Pointer KB, Clark PA, Eliceiri KW, Salamat MS, Robertson GA, Kuo JS. Administration of Non-Torsadogenic human Ether-à-go-go-Related Gene Inhibitors Is Associated with Better Survival for High hERG–Expressing Glioblastoma Patients. Clinical Cancer Research. 2017;23:73–80.

35. Besson P, Driffort V, Bon É, Gradek F, Chevalier S, Roger S. How do voltage-gated sodium channels enhance migration and invasiveness in cancer cells? Biochimica et Biophysica Acta (BBA) - Biomembranes. 2015;1848:2493–501.

36. Phan NN, Wang C-Y, Chen C-F, Sun Z, Lai M-D, Lin Y-C. Voltage-gated calcium channels: Novel targets for cancer therapy. Oncology Letters. 2017;14:2059–74.

37. Madden SK, de Araujo AD, Gerhardt M, Fairlie DP, Mason JM. Taking the Myc out of cancer: toward therapeutic strategies to directly inhibit c-Myc. Molecular Cancer. 2021;20:3.

38. Chen H, Liu H, Qing G. Targeting oncogenic Myc as a strategy for cancer treatment. Signal Transduction and Targeted Therapy. 2018;3:5.

39. Smith JC, Sheltzer JM. Genome-wide identification and analysis of prognostic features in human cancers. Cell Reports. 2022;38:110569.

40. Manville RW, Abbott GW. In silico re-engineering of a neurotransmitter to activate KCNQ potassium channels in an isoform-specific manner. Communications Biology. Nature Publishing Group; 2019;2:401.

41. Fukushiro-Lopes DF, Hegel AD, Rao V, Wyatt D, Baker A, Breuer E-K, et al. Preclinical study of a Kv11.1 potassium channel activator as antineoplastic approach for breast cancer. Oncotarget. Impact Journals, LLC; 2018;9:3321–37.

42. Gao J, Aksoy BA, Dogrusoz U, Dresdner G, Gross B, Sumer SO, et al. Integrative analysis of complex cancer genomics and clinical profiles using the cBioPortal. Science Signaling. 2013;

43. Gu Z, Eils R, Schlesner M. Complex heatmaps reveal patterns and correlations in multidimensional genomic data. Bioinformatics. 2016;32:2847–9.

44. Canisius S, Martens JWMM, Wessels LFAA. A novel independence test for somatic alterations in cancer shows that biology drives mutual exclusivity but chance explains most co-occurrence. Genome Biology [Internet]. BioMed Central; 2016 [cited 2020 Jan 17];17:261. Available from: http://genomebiology.biomedcentral.com/articles/10.1186/s13059-016-1114-x

45. Gel B, Serra E. karyoploteR: an R/Bioconductor package to plot customizable genomes displaying arbitrary data. Bioinformatics. 2017;33:3088–90.

46. Martincorena I, Raine KM, Gerstung M, Dawson KJ, Haase K, van Loo P, et al. Universal Patterns of Selection in Cancer and Somatic Tissues. Cell. 2017;171:1029–1041.e21.

47. Quinlan AR, Hall IM. BEDTools: A flexible suite of utilities for comparing genomic features. Bioinformatics. 2010;

48. Rosenthal R, McGranahan N, Herrero J, Taylor BS, Swanton C. deconstructSigs: Delineating mutational processes in single tumors distinguishes DNA repair deficiencies and patterns of carcinoma evolution. Genome Biology. 2016;

49. Davidson-Pilon C. lifelines: survival analysis in Python. Journal of Open Source Software. 2019;4.

50. Therneau TM, Lumley T. Survival Analysis; [R package “survival” version 3.1-12]. Comprehensive R Archive Network (CRAN). 2020;2.

51. Sun J, MacKinnon R. Cryo-EM Structure of a KCNQ1/CaM Complex Reveals Insights into Congenital Long QT Syndrome. Cell. 2017;

52. Edgar RC. MUSCLE: Multiple sequence alignment with high accuracy and high throughput. Nucleic Acids Research. 2004;

53. Feyfant E, Sali A, Fiser A. Modeling mutations in protein structures. Protein Science. 2007;

54. Abraham MJ, Murtola T, Schulz R, Páll S, Smith JC, Hess B, et al. GROMACS: High performance molecular simulations through multi-level parallelism from laptops to supercomputers. SoftwareX. Elsevier; 2015;1–2:19–25.

55. Lee S, Tran A, Allsopp M, Lim JB, Hénin J, Klauda JB. CHARMM36 united atom chain model for lipids and surfactants. Journal of Physical Chemistry B. 2014;

56. Stansfeld PJ, Goose JE, Caffrey M, Carpenter EP, Parker JL, Newstead S, et al. MemProtMD: Automated Insertion of Membrane Protein Structures into Explicit Lipid Membranes. Structure. 2015;

57. Monticelli L, Kandasamy SK, Periole X, Larson RG, Tieleman DP, Marrink S-J. The MARTINI Coarse-Grained Force Field: Extension to Proteins. Journal of Chemical Theory and Computation. American Chemical Society; 2008;4:819–34.

58. Stansfeld PJ, Sansom MSPP. From Coarse Grained to Atomistic: A Serial Multiscale Approach to Membrane Protein Simulations. Journal of Chemical Theory and Computation. American Chemical Society; 2011;7:1157–66.

59. Hall BA, Halim KBA, Buyan A, Emmanouil B, Sansom MSP. Sidekick for Membrane Simulations: Automated Ensemble Molecular Dynamics Simulations of Transmembrane Helices. Journal of Chemical Theory and Computation. American Chemical Society; 2014;10:2165–75.

60. Smart OS, Neduvelil JG, Wang X, Wallace BA, Sansom MSP. HOLE: A program for the analysis of the pore dimensions of ion channel structural models. Journal of Molecular Graphics. 1996;

61. Humphrey W, Dalke A, Schulten K. VMD: Visual molecular dynamics. J Mol Graph. 1996;14:33–8, 27–8.

62. Subramanian A, Tamayo P, Mootha VK, Mukherjee S, Ebert BL, Gillette MA, et al. Gene set enrichment analysis: A knowledge-based approach for interpreting genome-wide expression profiles. Proc Natl Acad Sci U S A. 2005;

63. Love MI, Huber W, Anders S. DESeq2. Genome Biol. 2014.

64. Aubrey BJ, Kelly GL, Kueh AJ, Brennan MS, O’Connor L, Milla L, et al. An inducible lentiviral guide RNA platform enables the identification of tumor-essential genes and tumor-promoting mutations in vivo. Cell Rep. 2015;10:1422–32.

65. Lou E, Fujisawa S, Morozov A, Barlas A, Romin Y, Dogan Y, et al. Tunneling nanotubes provide a unique conduit for intercellular transfer of cellular contents in human malignant pleural mesothelioma. PLoS One. 2012;7:e33093.

66. Yeung SYM, Greenwood IA. Electrophysiological and functional effects of the KCNQ channel blocker XE991 on murine portal vein smooth muscle cells. Br J Pharmacol. 2005;146:585–95.

67. Punke MA, Friederich P. Amitriptyline is a potent blocker of human Kv1.1 and Kv7.2/7.3 channels. Anesth Analg. 2007;104:1256–64, tables of contents.

68. Stirling DR, Carpenter AE, Cimini BA. CellProfiler Analyst 3.0: accessible data exploration and machine learning for image analysis. Bioinformatics. 2021;37:3992–4.

