## Supplementary Figures for "THE ROLE OF POTASSIUM CHANNELS IN THE PATHOGENESIS OF GASTROINTESTINAL CANCERS AND THERAPEUTIC POTENTIAL"

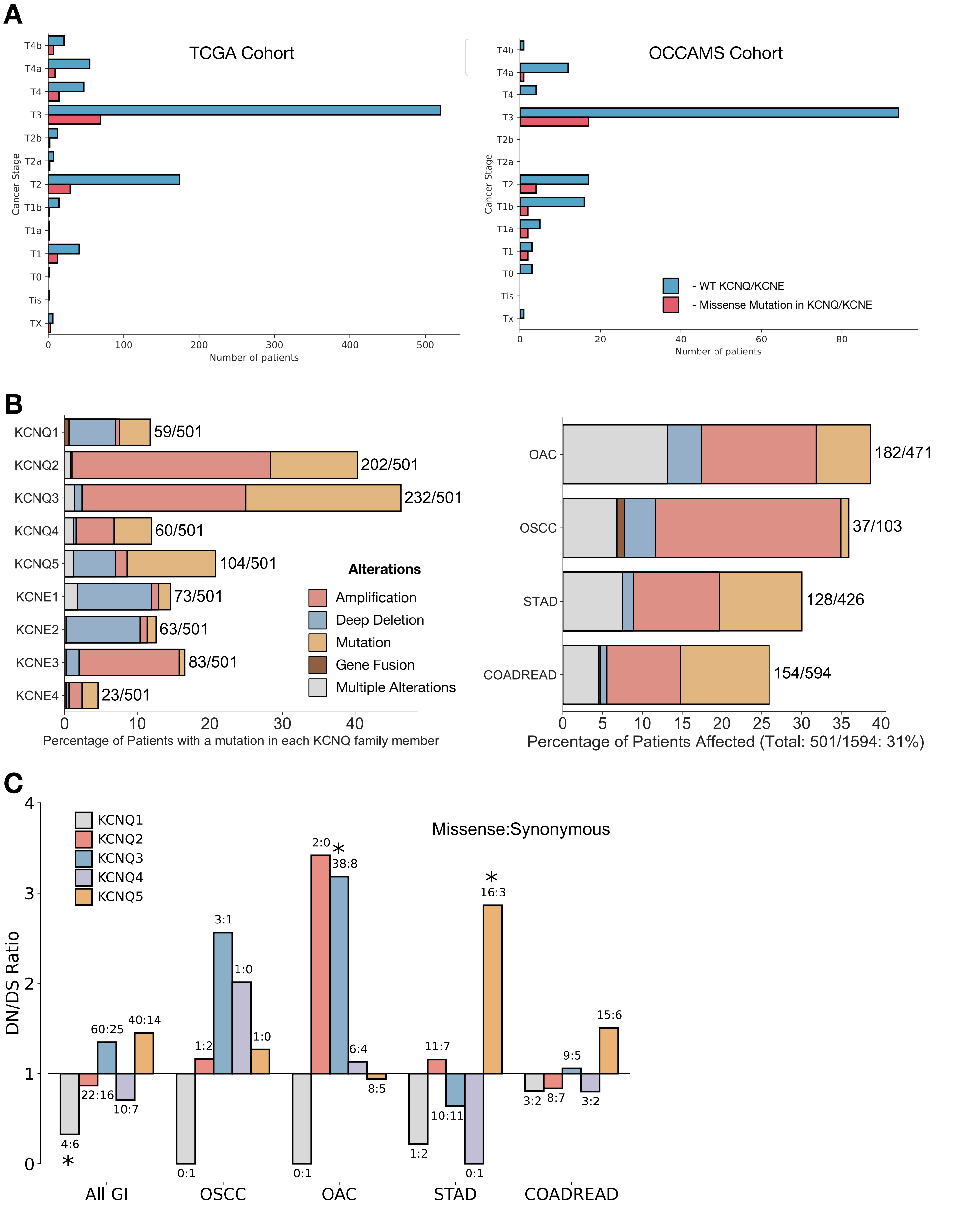


### Figure S1: ***KCNQ* genes are highly genetically altered in GI cancers, related to Figure 1**

A) Missense mutations in KCNQ genes in GI cancers by tumor stage

B) Percentage of types of genetic alterations in KCNQ/KCNE genes per patient (left) and per tissue (right)

C) dN/dS values for KCNQ genes in GI cancers.

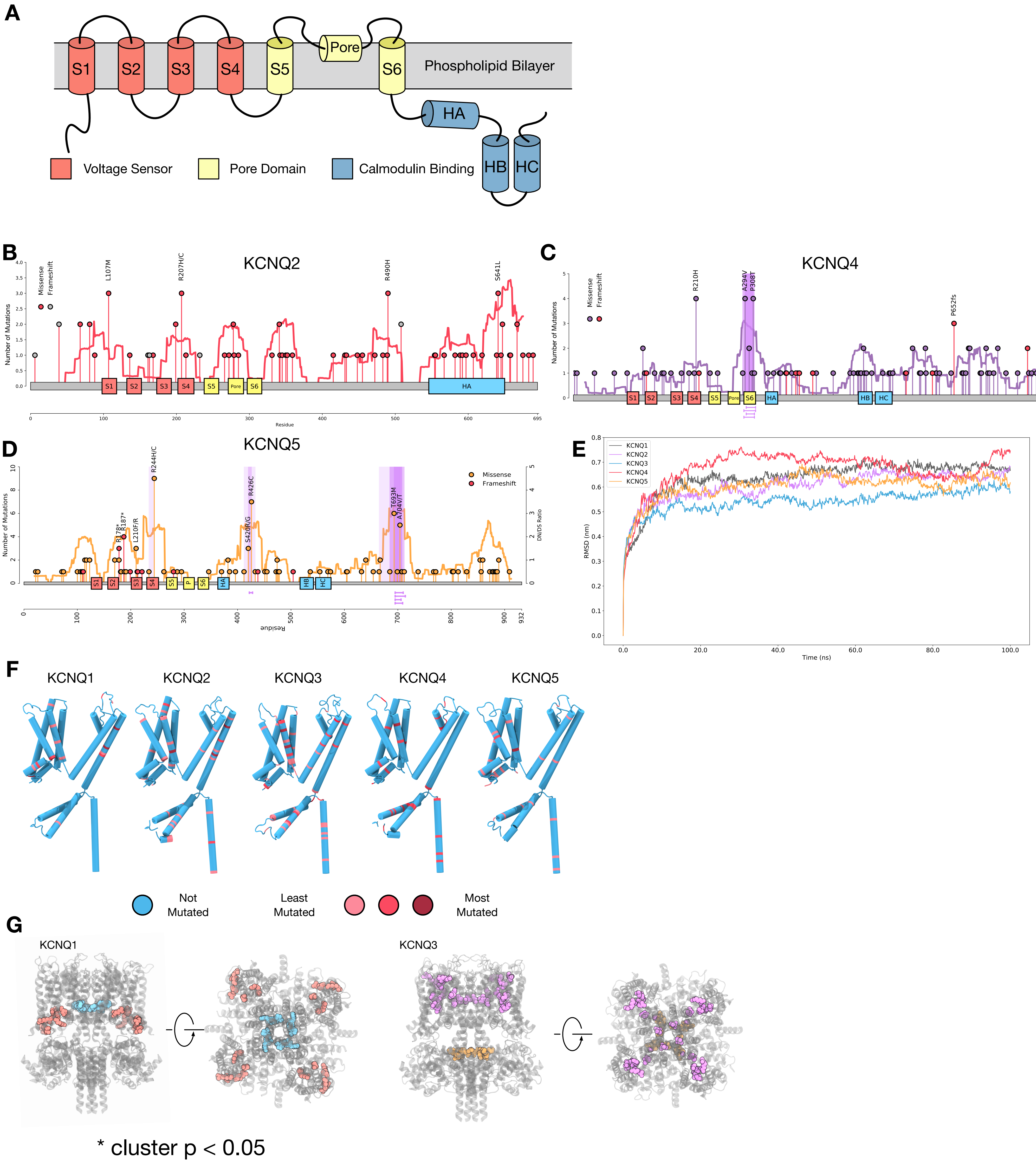


**Figure S2:** **Mutations in *KCNQ* genes are under selection, related to Figure 2**

A) Schematic of the KCNQ protein topology.

Mutational lollipop plots for: B) KCNQ2, C) KCNQ4, D) KCNQ5, purple highlights represent significant clusters ($q < 0.05$) as defined by the NMC algorithm.

E) Root Mean Square Deviation (RMSD) plots for simulations of KCNQ1-5 homology models in a POPC bilayer using 100ns of atomistic molecular dynamics.

F) Structure of single subunit of KCNQ1-5 colours by mutational frequency in GI cancer.

G) Structure and 3D clusters in KCNQ1 (left) and KCNQ3 (right), p value determined through permutation-based statistical test (see methods).

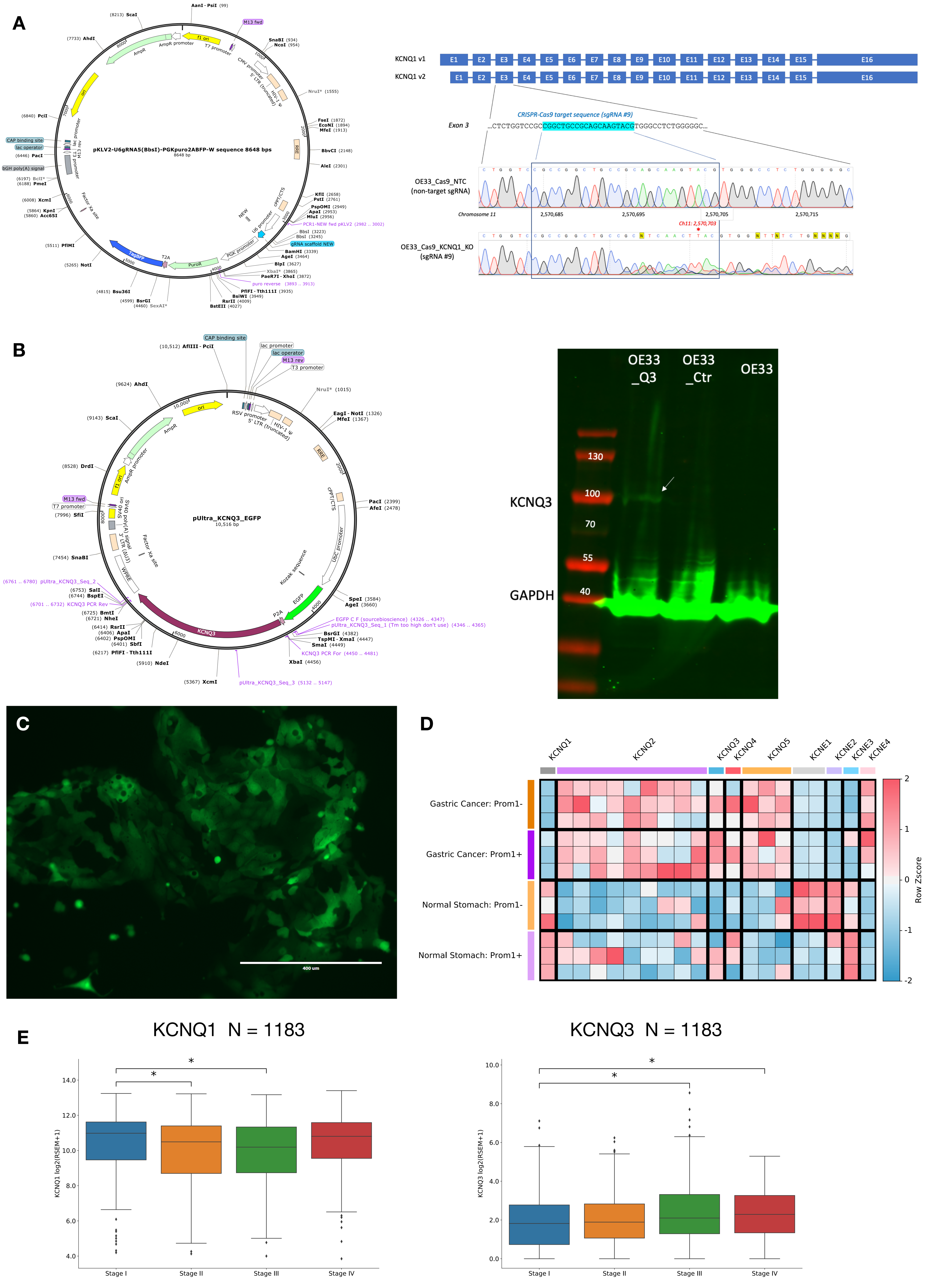


**Figure S3:** **Manipulation of KCNQ expression in vitro, and assessment of KCNQ expression in mouse models, related to Figure 3**

A) Plasmid used for KCNQ1 KO (left) Sanger sequencing confirming KCNQ1 KO in OE33 cell lines (right)

B) Plasmid used for KCNQ3 overexpression (left) Western blot validation of KCNQ3 expression in OE33 (right).

C) Confocal microscopy of KCNQ3 OE OE33 showing presence of GFP tag.

D) Microarray expression levels of KCNQ and KCNE genes in stomach cancer mouse model.

E) Expression of KCNQ1 (left) and KCNQ3 (right) in patients by tumor stage (AJCC tumor staging). * p represents students t-test < 0.05.

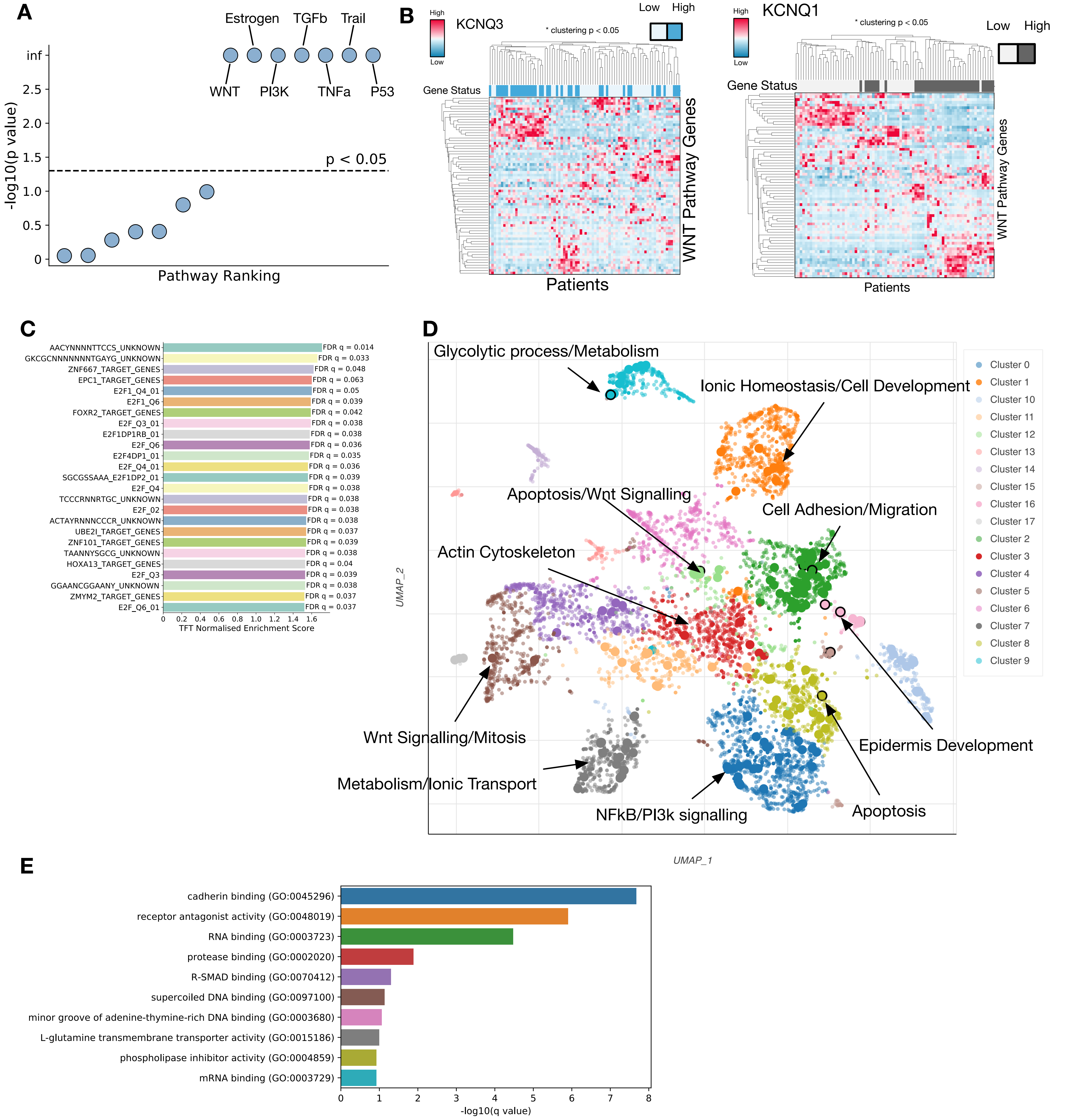


**Figure S4:** **KCNQ gene expression impacts WNT signalling, related to Figure 4**

A) -log10(pvalue) for PROGENY pathways correlated against KCNQ1 expression in human GI cancers.

B) Clustering of top and bottom KCNQ3 (left) and KCNQ1 (right) 25 expressing patients with GI cancers by WNT pathway genes. * p represents permutation clustering test (see methods)

C) Transcription factor enrichment scores for KCNQ3 OE vs WT OE33. Enrichment determined by GSEA against the TFT gene set.

D) Top GO Biological Processes enriched for differentially expressed genes in KCNQ3 OE vs WT OE33.

E) Top GO Molecular Function enriched pathways for differentially expressed genes in KCNQ3 OE vs WT OE33.

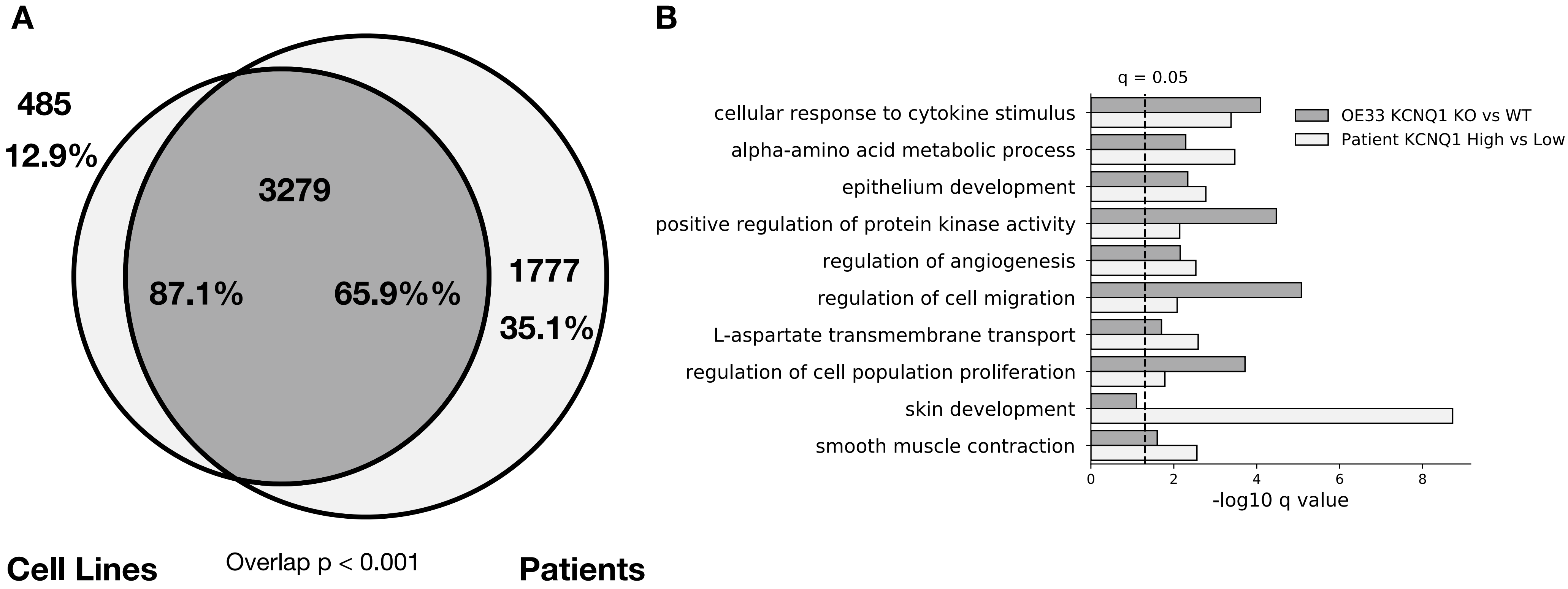


**Figure S5: OE33 cell lines moduled for KCNQ1 accurately reflect patients, related to figure 5.**

A) Venn diagram of overlap between enriched pathways in cell lines (KCNQ1 WT vs KCNQ1 KO OE33), and patients (highest 25 vs lowest 25 patients by KCNQ1 expression in OAC). Overlap p represents using cell line pathways as custom set in g-proflier.

B) -log10 q values for the top 10 overlapping pathways between cell lines and patients.

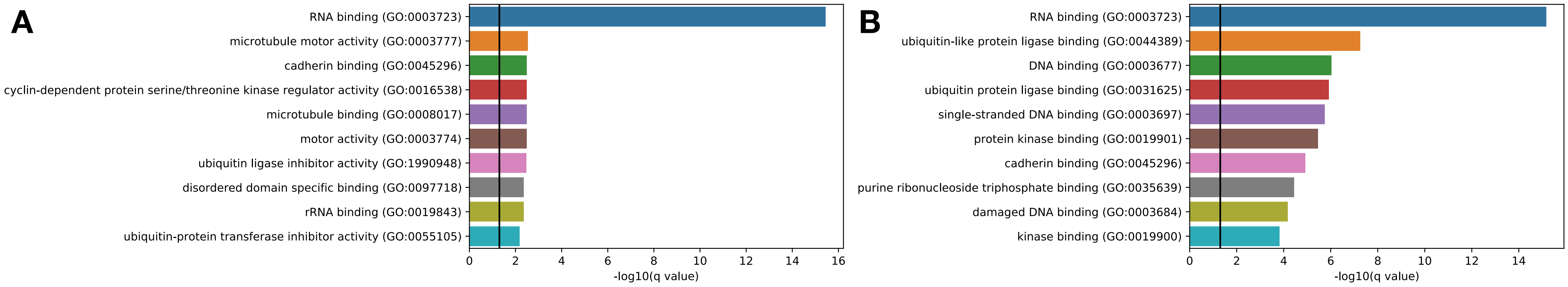


**Figure S6: Pathway enrichment for Linopirdine and Amitriptyline treated KCNQ3 OE OE33. Related to Figure 6.**

A) GO Molecular Function -log10(pvalues) for genes differentially expressed in ctrl vs Linopirdine treated KCNQ3 OE OE33.

B) GO Molecular Function -log10(pvalues) for genes differentially expressed in ctrl vs Amitriptyline treated KCNQ3 OE OE33.

#### Supplementary Tables:

Table S1: Mutual Exclusivity and Co-occurence values for KCNQ/E genes and a set of known GI cancer driver genes.

Table S2: Mutations with known functional/clinical consequence in the KCNQ family that are also observed in GI cancer.

Table S3: GSEA pathway analysis on the Hallmarks gene set for WT OE33 compared to KCNQ3 OE OE33.

Table S4: GSEA pathway analysis on the GO:Biological Processes gene set for WT OE33 compared to KCNQ3 OE OE33.

Table S5: Differential expression analysis results for WT OE33 compared to KCNQ3 OE OE33.

Table S6: Top enriched GO:Biological Processes for differentially expressed genes (padj < 0.05) in WT OE33 vs KCNQ3 OE OE33 cell lines.

Table S7: Top enriched GO:Biological Processes for differentially expressed genes (padj < 0.05) in top and bottom 25 expressers of KCNQ3 in GI cancer.

Table S8: Top enriched GO:Biological Processes for differentially expressed genes (padj < 0.05) in WT OE33 vs KCNQ1 KO OE33 cell lines.

Table S9: Top enriched GO:Biological Processes for differentially expressed genes (padj < 0.05) in top and bottom 25 expressers of KCNQ1 in GI cancer.

Table S10: Differentially expressed genes for KCNQ3 OE OE33 control vs 100mg/ml linopirdine.

Table S11: Differentially expressed genes for KCNQ3 OE OE33 control vs 100mg/ml amitriptyline.

Table S12: REVIGO GO:Biological Processes pathway clusters for pathways shared between 100mg/ml linopirdine and 100mg/ml amitriptyline exposure in KCNQ3 OE OE33.
